# *BiocPkgTools*: Toolkit for Mining the *Bioconductor* Package Ecosystem

**DOI:** 10.1101/642132

**Authors:** Shian Su, Vincent J. Carey, Lori Shepherd, Matthew Ritchie, Martin T. Morgan, Sean Davis

## Abstract

**Motivation:** The *Bioconductor* project, a large collection of open source software for the comprehension of large-scale biological data, continues to grow with new packages added each week, motivating the development of software tools focused on exposing package metadata to developers and users. The resulting BiocPkgTools package facilitates access to extensive metadata in computable form covering the *Bioconductor* package ecosystem, facilitating downstream applications such as custom reporting, data and text mining of *Bioconductor* package text descriptions, graph analytics over package dependencies, and custom search approaches.

**Results:** The BiocPkgTools package has been incorporated into the *Bioconductor* project, installs using standard procedures, and runs on any system supporting R. It provides functions to load detailed package metadata, longitudinal package download statistics, package dependencies, and *Bioconductor* build reports, all in “tidy data” form. BiocPkgTools can convert from tidy data structures to graph structures, enabling graphbased analytics and visualization. An end-user-friendly graphical package explorer aids in task-centric package discovery. Full documentation and example use cases are included.

**Availability:** The BiocPkgTools software and complete documentation are available from *Bioconductor* (https://bioconductor.org/packages/BiocPkgTools).

## Introduction

*Bioconductor* is a open source software project (comprising 1741 individual analysis packages) and community for the analysis and comprehension of large-scale biological data. Newly submitted software packages undergo a technical review to ensure that best practices and *Biocondutor* coding conventions are followed. The project maintains an automated build system that ensures that packages in the *Bioconductor* project are compiled and built successfully and pass basic checks. Package downloads are tracked and aggregated by package and month, longitudinally. Finally, package details such as title, description, version, author, and dependencies on other R packages are compiled based on package metadata.

The current size and growth of the *Bioconductor* project suggests that there is merit in exposing computable forms of the metadata describing the *Bioconductor* package ecosystem. To that end, we developed a small suite of tools to provide easy access to project details such as download statistics, bulk package metadata, and package build status. Developers, project leaders, open source software researchers, and *Bioconductor* end users can build on the availability of these data for applications such as custom reporting, dependency graph analytics, package filtering, and text mining.

### Features and usage

The core functionality of BiocPkgTools is to expose *Bioconductor* project and package metadata as tidy data [3] objects. The data presented by the package are accessed directly from online resources available from *Bioconductor*. As such, the package relies on web connectivity and collects the most recent data. Installation instructions are detailed on the package website.

Package functionality can be roughly divided into data access, data presentation, and graph/network functionality. See Table 1 for an overview.

**Table 1:** Main package functions and descriptions.

| Name | Functionality |
| --- | --- |
| biocPkgList | Package details including description, author and maintainer, dependencies, URLs, bug report mechanism |
| biocDownloadStats | Monthly download statistics for all packages |
| biocbuildReport | *Bioconductor* build report for all packages and systems |
| biocExplore | Interactive, browsable “bubble plot” of *Bioconductor* packages and details |
| problemPage | Interactive, customized build report for an individual package author |
| buildPkgDependencyDataFrame | Package dependencies as data frame |
| buildPkgDependencyIgraph | Package dependencies as a graph [2] |
| inducedSubgraphByPkgs | Create a minimal subgraph of *Bioconductor* dependencies based on specific packages |
| subgraphByDegree | Create a subgraph of all packages within a given degree of a single package |

After installing BiocPkgTools, the `biocDownloadStats` function can generate a tidy data structure summarizing monthly download statistics (both total and unique IP addresses) for all *Bioconductor* packages.

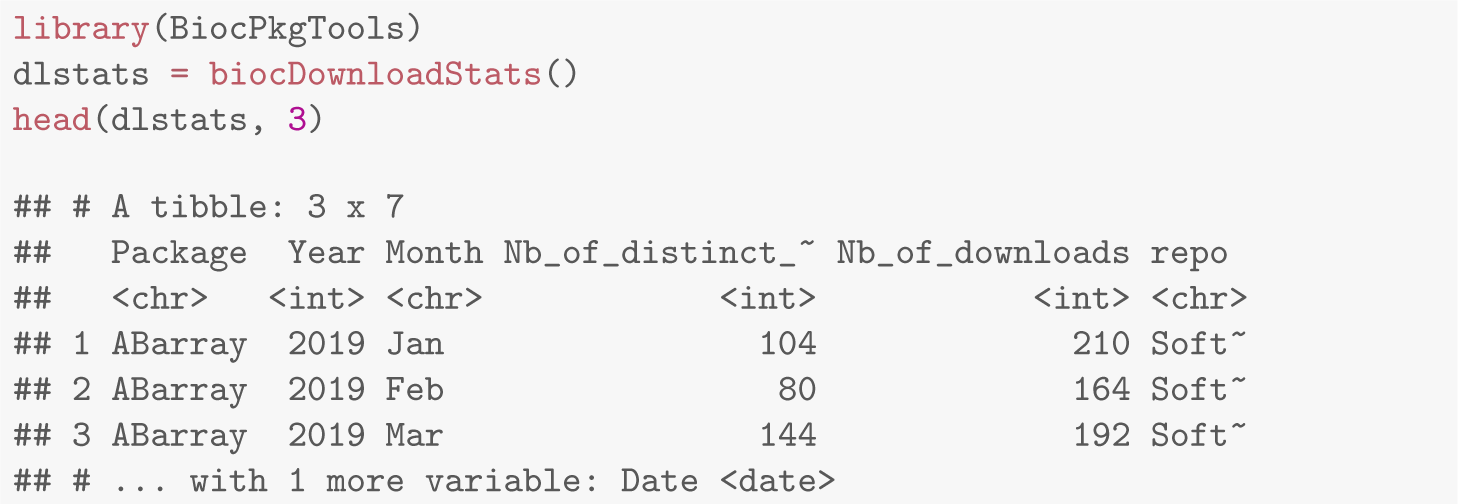

The `biocBuildReport` function gathers information from the *Bioconductor* build report site *and can be used, for example, to summarize the “build status” for all Bioconductor* pacakages.

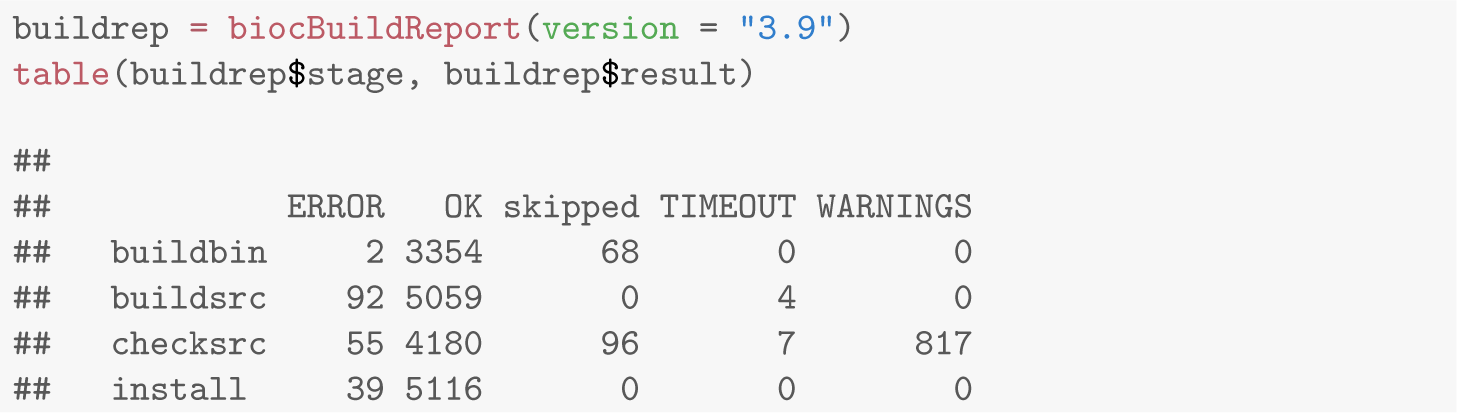

These data are useful to developers to track the health of their software either programmatically or via a searchable, sortable table from the `problemPage` function.

As an alternative to basic web browser search and the *Bioconductor* online software list, the `biocExplore` function offers interactive and graphical approach to package browsing (see Figure 2). The biocExplore widget allows browsing packages under different biocViews, Bioconductor’s software catergory tags. This interactively visualises the relative number of downloads each package has under different biocViews, allowing users to quickly determine which packages are most commonly used for different analysis tasks.

**Figure 1:**
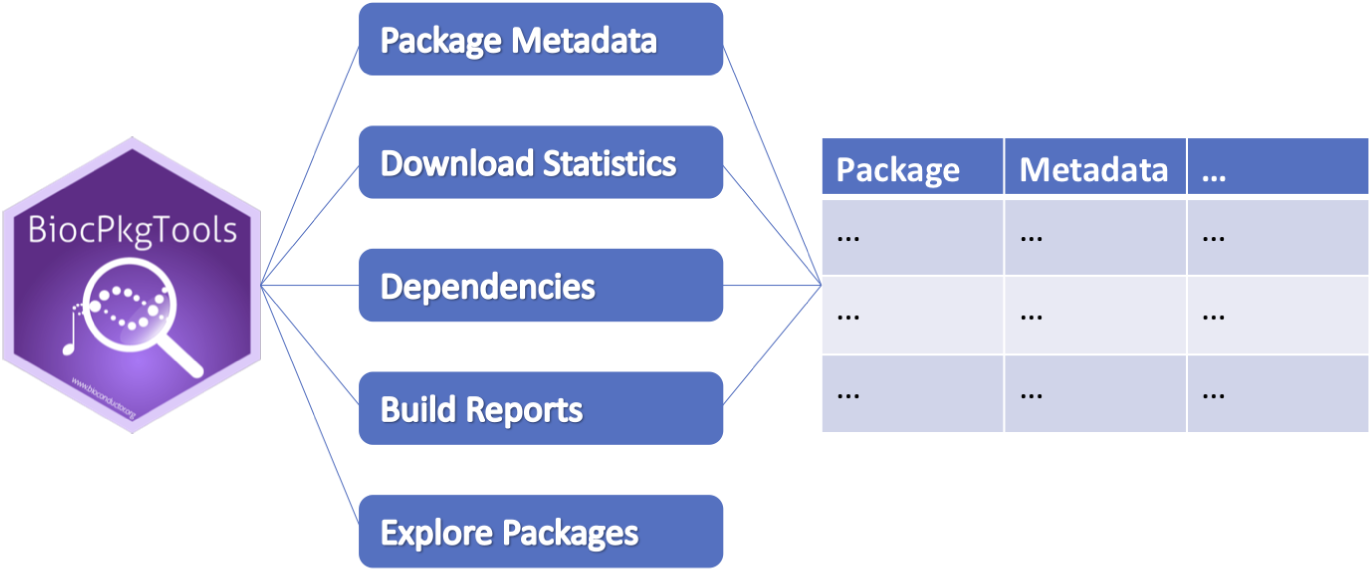
Schematic overview of the BiocPkgtools package. BiocPkgTools can access and transform web-accessible resources including package metadata, download statistics, dependencies between packages, and updated *Bioconductor* build report status to “tidy data” reports that can be manipulated using standard R tools. Interactive package exploration is also available.

**Figure 2:**
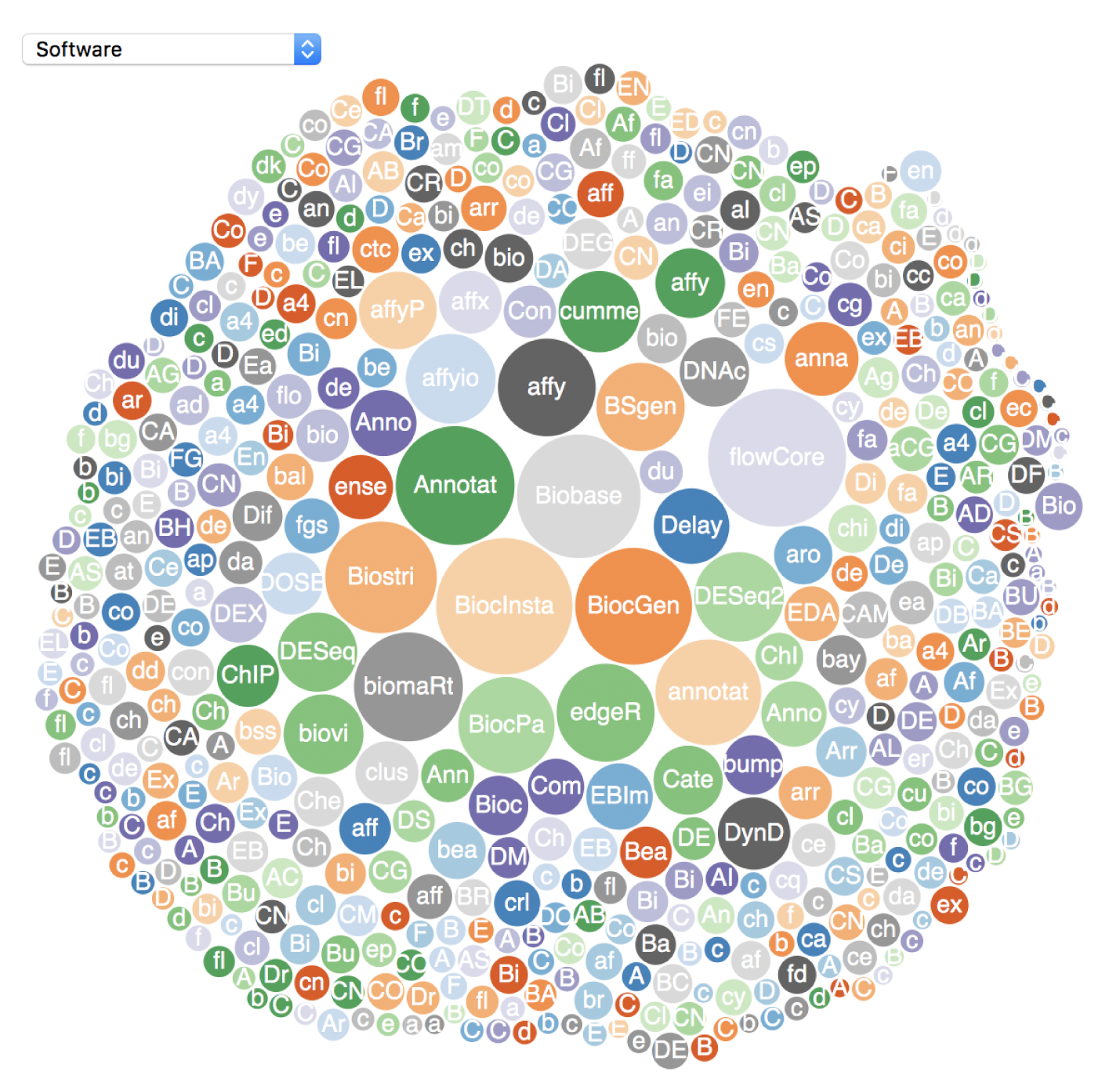
The `biocExplore` function opens an interactive web application that allows users to select focused groups of *Bioconductor* packages to view as a bubble plot. Bubbles are sized based on download statistics. Hovering over a bubble will give download number while clicking on a bubble will pop up a package details page, including a link to the package landing page.

The *Bioconductor* package ecosystem is, by design, highly interconnected via package dependencies. Several functions in the BiocPkgTools package provide examples of package dependency graph creation and visualization. Figure 3 displays packages within one degree of dependency relationship of the GEOquery package.

**Figure 3:**
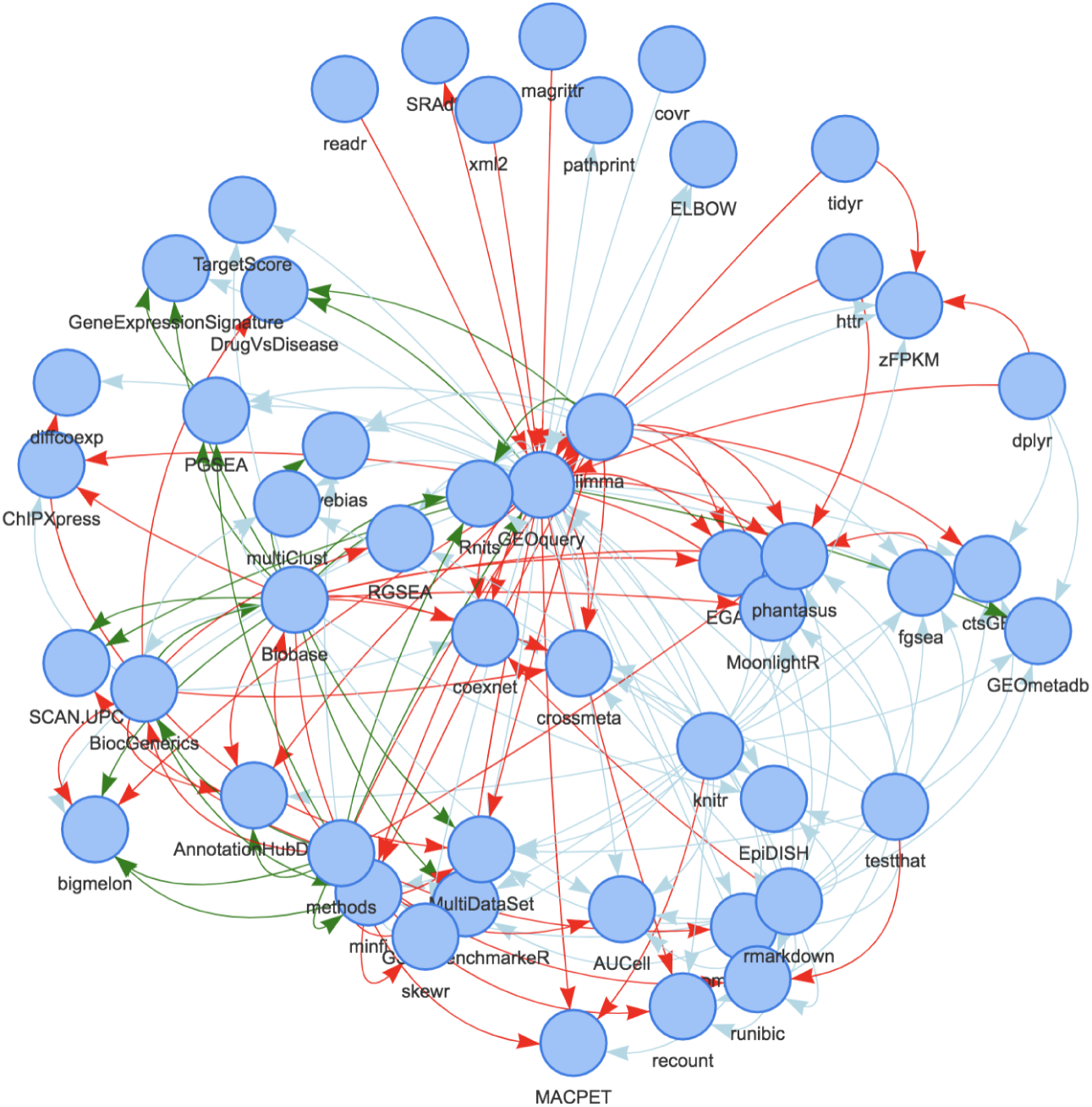
The `subgraphByDegree` function builds a data visualization of dependencies between all packages within one degree of the GEOquery package using the visNetwork package [1]. Links are colored based on type (Suggests [light blue], Depends [green], and Imports [red]) and arrows point to the “dependent” package.

## Discussion

The BiocPkgTools package comprises a set of functions for accessing software metadata from the growing collection of *Bioconductor* packages. For software developers, this metadata can be useful for tracking package build status and the health of package dependencies. Easy access to descriptive package metadata for all *Bioconductor* software resources can empower researchers or users interested in text mining, custom package search, or analysis of the existing software ecosystem. BiocPkgTools can provide easy access to metrics of *Bioconductor* sofware usage that are increasingly being incorporated into funding and promotion decisions.

## Implementation

BiocPkgTools is implemented as a standard R package and hosted in the *Bioconductor* repository. All functions are documented and include examples. An included tutorial (vignette) demonstrates features and capabilities.

## Data availability

analyses are included All data accessed and used by the BiocPkgTools package are publicly available and are updated regularly at the *Bioconductor* project.

## Software availability

This section will be generated by the Editorial Office before publication. Authors are asked to provide some initial information to assist the Editorial Office, as detailed below.

- Software URL: https://bioconductorg.org/packages/BiocPkgTools
- Version Control URL: https://github.com/seandavi/BiocPkgTools
- Software package DOI: doi:10.18129/B9.bioc.BiocPkgTools
- Software License: MIT License

## Author contributions

SD conceived of the project. All authors have reviewed and edited the manuscript. Authors contributing to code include MM, SD, SS, LS, and VC.

